# Cannabidiol attenuates cognitive deficits and neuroinflammation induced by early alcohol exposure in a mice model

**DOI:** 10.1101/2021.03.16.435465

**Authors:** Alba García-Baos, Xavi Puig-Reyne, Óscar García-Algar, Olga Valverde

## Abstract

Foetal alcohol spectrum disorder (FASD) is the umbrella term used to describe the physical and mental disabilities induced by alcohol exposure during development. Early alcohol exposure induces cognitive impairments resulting from damage to the central nervous system (CNS). The neuroinflammatory response accompanied by neurodegenerative mechanisms contribute to those detrimental alterations. Cannabidiol (CBD) has recently emerged as an anti-inflammatory drug that might be useful to treat several neuropsychiatric disorders. In our study, we assessed the effects of CBD on long-lasting cognitive deficits induced by early alcohol exposure. Furthermore, we analysed long-term pro-inflammatory and apoptotic markers within the prefrontal cortex and hippocampus. To model alcohol binge drinking during gestational and lactation periods, we used pregnant C57BL/6 female mice with time-limited access to 20% v/v alcohol solution. Following the prenatal and lactation alcohol exposure (PLAE), we treated the male and female offspring with CBD from post-natal day (PD) 25 until PD34, and we evaluated their cognitive performance at PD60. Our results showed that CBD treatment during peri-adolescence period ameliorates cognitive deficits observed in our FASD-like mouse model, without sex differences. Moreover, CBD restores the PLAE-induced increased levels of TNFα and IL-6 in the hippocampus. Thus, our study provides new insights for CBD as a therapeutic agent to counteract cognitive impairments and neuroinflammation caused by early alcohol exposure.

## 1. INTRODUCTION

Alcohol is alarmingly used by pregnant women nowadays, and approximately one third of those who reported alcohol consumption over pregnancy engaged in binge drinking [1], which is particularly harmful to the brain of a developing foetus [2]. FASD comprises a wide range of morphological and functional anomalies that arise from early alcohol exposure. The consequences of alcohol drinking during pregnancy and/or breastfeeding may vary in severity across the spectrum [3]. However, cognitive impairments are core symptoms among FASD patients, especially learning and memory deficits which are highly incident in all diagnoses [4].

Since FASD is a neurodevelopmental disorder, an early intervention is key to minimize the devastating effects of alcohol exposure. Despite the lack of curative treatment, miscellaneous interventions have been employed to ameliorate their quality of life, such as parental training, behavioural education, and pharmacotherapy. Thereby, anti-inflammatory agents have been proposed as treatment for FASD to address the neuroinflammatory phenotype observed in such patients [5,6]. In fact, previous studies showed that prenatal alcohol exposure induces microglial activation as well as increased production of cytokines in the foetal rodent brain [7–9], which might persist until the adulthood [10]. Importantly, earlier preclinical research indicated that deficiency of toll-like receptor 4 (TLR4), a receptor involved in the activation of innate immune response [11], prevents the pro-inflammatory response along with long-term behavioural impairments found in alcohol exposed offspring [12]. In addition, the neuroinflammatory and apoptotic signalling in cerebral cortex and hippocampus can be blocked by the anti-inflammatory agent resveratrol, preventing cognitive deficits in early alcohol exposed rats [13]. Altogether, these findings support that the neuroinflammatory state induced by developmental alcohol exposure is a relevant trigger for long-term cognitive dysfunctions.

In the last decades, studies have shown that some cannabinoids have a neuroprotective pharmacological profile through multiple mechanisms [14]. These compounds exhibit a potential therapeutic role in the context of neuroinflammatory and neurodegenerative disorders, including the alcohol-induced neuroinflammation [15]. Among phytocannabinoids, cannabidiol (CBD) is able to interact with multiple targets within the central nervous system (CNS) exerting distinct pharmacological effects [14]. For instance, CBD increases the levels of endocannabinoids as consequence of fatty acid amide hydrolase enzyme (FAAH) inhibition [16]. The increased tone of endocannabinoids decreases neuronal damage, which is prevented by cannabinoid receptor type 1 (CB1R) and cannabinoid receptor type 2 (CB2R) antagonisms [17]. CBD can activate non-cannabinoid receptor pathways to exert its neuroprotective effects as well [18]. Previous studies showed that CBD can reduce microglial activation and pro-inflammatory cytokine production [19,20]. Although the mechanisms underlying its anti-inflammatory effects are not entirely understood, the activation of peroxisome proliferator-activated receptor (PPAR) gamma [21,22] could play a key role. Other receptors, such as transient receptor potential vanilloid 1 [23,24] or G protein-coupled receptor 55, might also be involved in these responses [18].

Additionally, CBD might be a promising therapeutic candidate for the management of sequels induced by excessive alcohol exposure. Preclinical data revealed that CBD attenuates the neurodegeneration within hippocampus [25] and entorhinal cortex [25,26] caused by a rodent model of binge-like alcohol drinking. Notwithstanding, CBD effect on developmental alcohol exposure has never been addressed.

To evaluate the impact of alcohol exposure during early developmental periods in the offspring, we have developed a reliable mouse model of binge-like alcohol drinking during gestational and lactation periods. Our previous findings indicated that early alcohol-exposed mice show spatial, working and recognition memory impairments [10,27], which mimics the behavioural impairments found in FASD patients. In the present study, we aimed to explore the therapeutic effects of CBD demonstrating that the phytocannabinoid might attenuate cognitive deficits induced by developmental alcohol exposure, conceivably through an anti-inflammatory mechanism.

## 2. MATERIALS AND METHODS

### 2.1. Animals

Male and female C57BL/6 were purchased from Charles River (Barcelona, Spain) and transported to our animal facility (UBIOMEX, PRBB) to be used as breeders. Animals were housed in a room with controlled temperature- (21 ± 1 °C), humidity- (55% ± 10%) and lighting (light remained on between 7:30 p.m. and 7:30 a.m.). Mice were allowed to acclimatize to the new environmental conditions for at least 1 week prior to experimentation, which occurred during the dark phase under a dim red light. Animals were 12 weeks old when the breeding began, and they were housed 1 male with 2 females. After successful mating, pregnant females were individualised and observed daily for parturition. For each litter, the date of birth was designated as PD0. Pups remained with their mothers for 21 days and were then weaned (PD21). After weaning, male and female offspring were housed in separated groups. Food and water were available *ad libitum*, except when water was substituted by alcohol to carry out the drinking in the dark (DID) test. All animal care and experimental procedures were conducted in accordance with the European Union Directive 2010/63/EU regulating animal research and were approved by the local Animal Ethics Committee (CEEA-PRBB).

### 2.2. DID test

This procedure was conducted as previously reported [10,27,28], which allows to model an early alcohol exposure under a binge-like drinking pattern. Briefly, two days after mating, pregnant females were randomly assigned to two groups: alcohol or water (control). Three hours after the lights were turned off, the water bottles were replaced with 10-ml graduated cylinders fitted with sipper tubes containing either 20% (v/v) alcohol in tap water or only tap water. From Monday to Wednesday (days 1, 2, 3), pregnant females were allowed to voluntarily drink for 2h-access period. On Thursday (day 4), the drinking-access period was extended to 4h. Volumes consumed were recorded just after the drinking-access periods. During this procedure, all the females were individually housed in order to record individual fluid intakes. Following the drinking-access period, water bottles were returned to the home cage. Fluid intakes (g/kg of body weight) were calculated for each day on the basis of average 2-day body weight values, since dams were weighed at 2-day intervals (Monday and Wednesday).

In our experiment, DID test consisted of six consecutive weeks, encompassing prenatal and lactation periods. Therefore, each pregnant dam was exposed to six binge-like drinking sessions (day 4), following the habituation period (days 1,2,3). As previously reported, blood alcohol concentration of dams reached levels of ~0.8 g/L after the last binge-like drinking session [10], which is in accordance with “Drinking Levels Defined” reported by NIAAA [29].

### 2.3. Drugs

Ethyl alcohol was purchased from Merck Chemicals (Darmstadt, Germany) and diluted in tap water to obtain a 20% (v/v) alcohol solution. CBD (20 mg/kg i.p.) was generously provided by Phytoplant Research S.L., (Córdoba, Spain). The dose of 20 mg/kg was chosen based on other studies showing this dose within the anti-inflammatory therapeutic range in rodents and humans [30]. CBD was first mixed with 2% Tween-80 and, after being homogenised by grinding in a mortar, 0.9% NaCl was added.

### 2.4. Experimental design

Offspring mice were administered with vehicle or CBD (20 mg/kg i.p.) during 10 consecutive days from PD25 until PD34, corresponding to the peri-adolescence period. The treatments were randomly assigned to either exposed to water (control) or to alcohol (PLAE group). Then, animals were left undisturbed at the home cage until reaching the adulthood (PD60), when the behavioural tests were carried out as shown in Figure 1A. Male and female offspring were used as a whole population since there were not differences due to the sex factor. All groups contained balanced number of male and female to avoid this possible bias. At least eight litters were used for each experimental group and treatments were randomly distributed through all litters to avoid the litter effect.

**Fig. 1.**
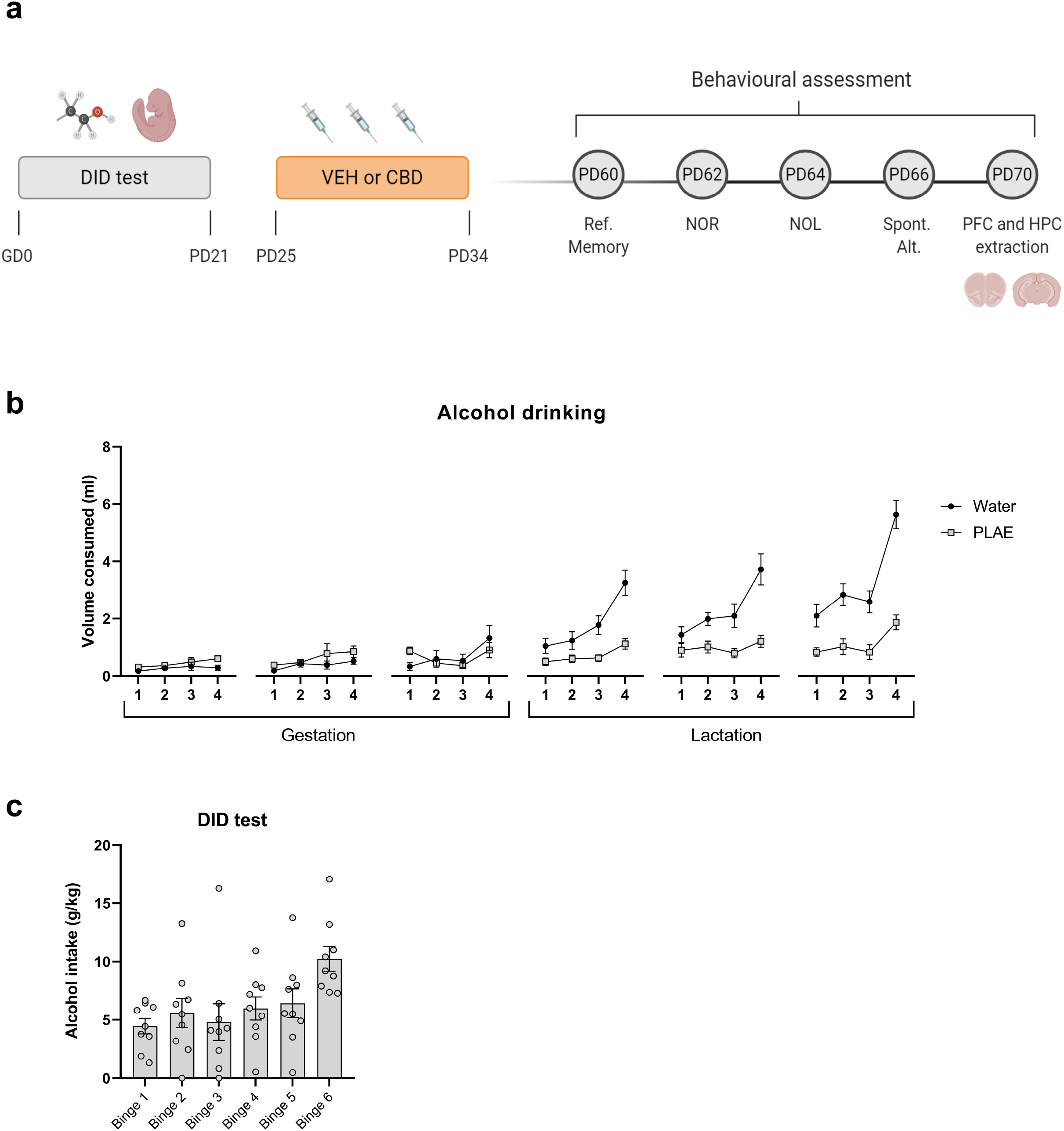
Experimental design and alcohol consumption by dams during binge-like drinking sessions. (a) Schematic representation of experimental design extensively described above. (b) Volumes (ml) of water or alcohol drinking by pregnant mice during DID test. (c) Alcohol intake (g of alcohol per kg of body weight) in the six binge-like drinking sessions throughout gestational and lactation periods (n=9 dams). Data are presented as mean ± SEM. CBD, cannabidiol; DID, drinking-in-dark test; HPC, hippocampus; NOL, novel object location; NOR, novel object recognition; PFC, prefrontal cortex; Ref. memory, reference memory; VEH, vehicle; Spont. Alt, spontaneous alternation

### 2.5. Reference memory test

A black Y-maze (two equal 395mm-long arms, separated by 120° angles) was employed to assess spatial reference memory [31]. This experiment consisted of two phases (training and testing session) with 1h of inter-trial interval, which is considered as short-term spatial memory. In the training session, one arm of the Y-maze was closed off, which was designated as the novel arm. Then, the mouse was placed into the start arm of the maze and was allowed to freely explore the two remaining arms for 5 minutes.

When the testing session started, the mouse was placed back into the maze with the blockage removed and was allowed to freely explore the three arms for 5 minutes. One visual clue was placed on the furthest wall of each arm in order to help the mouse recognize the arms of the maze. The time spent in each of the three arms was measured by the Smart Software (Panlab s.l.u., Barcelona, Spain). The preference ratio was calculated by: 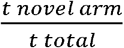 being “t” the time each mouse spent exploring the arms.

### 2.6. Novel Object Recognition (NOR) task

48h following the reference memory test, this task was performed as described previously [32] with minor modifications. Black open boxes (24 cm x 24 cm x 15 cm) and plastic toys of similar size to mice were used in this experiment. Briefly, the procedure consisted of three phases: habituation, training and test session, always under dim light intensity (30lux). During the habituation, mice were individually acclimatised to the box for 10 minutes without objects. After 2h, the training session took place, and the mice were allowed to explore the box in the presence of two identical objects (familiar objects) for 10 minutes. Finally, the test session was performed 4h after the training. In the test session, a familiar object was replaced by a novel one and the animals were allowed to explore for 10 minutes. The familiar and novel objects were counterbalanced. The discrimination index was calculated as: 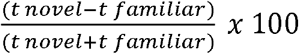, being “t” the time a mouse spent exploring the objects (manually recorded from a video by an observer who was blind to the experimental groups).

### 2.7. Novel Object Location (NOL) task

This task was performed 48h after NOR. We used black open boxes (24 cm x 24 cm x 15 cm) and two identical plastic toys, which were different from the ones used in the NOR task. The procedure consisted of the identic three phases as in the NOR task, under the same timing and lighting conditions. However, during the test session, one of the familiar objects was displaced to a new location on the arena. This change of spatial configuration triggers an increase level of exploration compared with the non-displaced object, as other authors have previously reported [33,34]. Here, the displaced and non-displaced objects were counterbalanced. In this case, the discrimination index was calculated as: 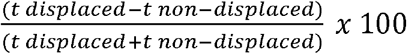, being “t” the time a mouse spent exploring the objects (manually recorded from a video by an observer who was blind to the experimental groups).

### 2.8. Delayed Spontaneous Alternation Task

This task is commonly used to assess spatial working memory [35,36]. The same Y-maze as in the section “2.5” was used to perform this task. Briefly, each mouse carried out 10 trials over one day, with inter-trial intervals of 30 minutes. Each trial was composed by two phases: the forced-choice and the free-choice phases, with an intra-trial delay of 40 seconds. First, in the forced-choice phase, mice were placed in the start arm of the Y-maze that had randomly one arm closed off, so they were forced to choose the other arm. Once the animal was completely inside the arm, it was closed for freely exploration for 10 seconds, and then removed from the maze. In the free-choice phase, just 40 seconds later, the animal was placed again in the start arm and it was allowed to freely choose the arm for exploration. A correct alternation was considered when the animal chose the non-forced arm in the free-choice phase. Then, the percentage of correct alternations was calculated considering 10 trials per mouse.

### 2.9. Western blotting assay

At PD70, animals were euthanized by cervical dislocation and PFC and HPC were dissected and stored at −80°C. The tissue was homogenized in lysis buffer [0.15M NaCl, 1% TX-100, 10% glycerol, 1mM EDTA, 50mM TRIS pH=7.4 and a phosphatase and protease inhibitor cocktail (Roche, Basel, Switzerland)], using 25 μl per mg of tissue. Homogenates were centrifuged at 1,000× g for 20 min at 4°C, and the resulting supernatants were collected for protein quantification. The lysate protein concentration was determined using a stock solution of 5 mg/ml BSA as a protein standard. Equal amount of protein (30 μg) for each sample were mixed with loading buffer (153 mM TRIS pH = 6.8, 7.5% SDS, 40% glycerol, 5 mM EDTA, 12.5% 2-β-mercaptoethanol, and 0.025% bromophenol blue), loaded onto 10% polyacrylamide gels, and then transferred to PVDF membranes (Immobilion-P, MERCK, Burlington, USA). Membranes were blocked for 30 minutes with 5% bovine serum albumin at RT and then immunoblotted overnight at 4°C using primary antibodies (see Table 1). On the next day, membranes were 3 times washed for 5 minutes each with T-TBS 1X [Tris-buffered saline (100 mmol/L NaCl, 10 mmol/L Tris, pH = 7.4) and 0.1% Tris-buffered saline Tween-20]. Then, membranes were incubated for 1h with their respective secondary fluorescent antibodies: anti-mouse (1:2,500, IRDye 800, Abcam, Cat# ab216772, RRID: AB_2857338) and anti-rabbit (1:2,500, DyLight™ 680, Rockland, Cat# 611-144-002, RRID: AB_1660962). Protein expression was quantified using an Amersham™ Typhoon™ scanner and quantified using Image Studio Lite software v5.2 (LICOR, USA). Protein expression signals were normalized to the detection of housekeeping control protein in the same samples and expressed in terms of fold-change with respect to control values.

**Table 1.** Primary antibodies

| Antibody | Description | Host | Dilution | Company |
| --- | --- | --- | --- | --- |
| **Caspase-3** | Cleaved caspase-3 (active form) | Rabbit | 1:1000 | Cell Signaling |
| **COX-2** | Cyclooxygenase-2 | Rabbit | 1:1000 | Cell Signaling |
| **IL-6** | Interleukin-6 | Mouse | 1:1000 | Santa Cruz<br>Biotechnology |
| **NLRP3** | NOD-like receptor family, pyrin domain-containing protein 3 | Rabbit | 1:1000 | GeneTex |
| **NFkB p65** | Nuclear factor Kappa-B p65 subunit (also known as RelA) | Mouse | 1:500 | Santa Cruz<br>Biotechnology |
| **TNFα** | Tumor necrosis factor alpha | Rabbit | 1:1000 | Cell Signaling |
| **Tubulin** | Class III β-Tubulin | Rabbit | 1:5000 | Abcam |
| **Tubulin** | β-Tubulin | Mouse | 1:5000 | BD Pharmigen |

### 2.10. Data and statistical analyses

Animals were randomly assigned to an experimental group. During the behavioural manipulations and data interpretation, researchers were blind to the treatment each animal had received. We analysed the results of all the behavioural test and western blotting analyses using two-way ANOVA with factors defined as *Group* (levels: Water and PLAE) and *Treatment* (levels: VEH and CBD). When *F* values achieved significance and there was no significant variance in homogeneity, Bonferroni’s *post hoc* tests were run. Pearson’s correlation analysis was used to explore correlations between the variables.

Data are presented as mean ± SEM. For statistical analyses we used GraphPad Prism 8.0. software. The α level of statistical significance was set at P < 0.05. The exact group size for the individual experiments is shown in the corresponding figure legends.

The schematic representation shown in Figure 1a was created in BioRender.com,

## 3. RESULTS

### 3.1. Maternal alcohol consumption

Dams were allowed to access either water or 20% alcohol following the 6 weeks of DID schedule explained above. Liquid volumes were recorded throughout the gestation and lactation periods. The water and alcohol consumption (volume in ml) over the 6 weeks is presented in Figure 1b. The fluid intake was then calculated only for dams exposed to alcohol and is represented in Figure 1c. Our team has previously demonstrated correlation between similar alcohol intakes and moderate-to-high blood alcohol concentration in dams [10] as well as in pups [27].

### 3.2. CBD treatment from PD25 to PD34 improves some cognitive deficits induced by PLAE

The four behavioural tasks were focused on the cognitive domain, specifically on memory functions.

#### 3.2.1. CBD counteracts the PLAE-induced effects on reference memory

The two-way ANOVA reveals a significant *interaction* effect between *group* x *treatment* factors (F_(1,45)_ = 4.906, p<0.05). Subsequent Bonferroni’s *post-hoc* analysis displays a tendency (p=0.06) to impair reference memory when comparing PLAE-VEH to the control group Water-VEH. Still, PLAE-CBD group showed significantly higher preference for the novel arm as compared to PLAE-VEH (p<0.05), revealing that CBD treatment hampers the detrimental effect on reference memory caused by PLAE (Figure 2a).

**Fig. 2.**
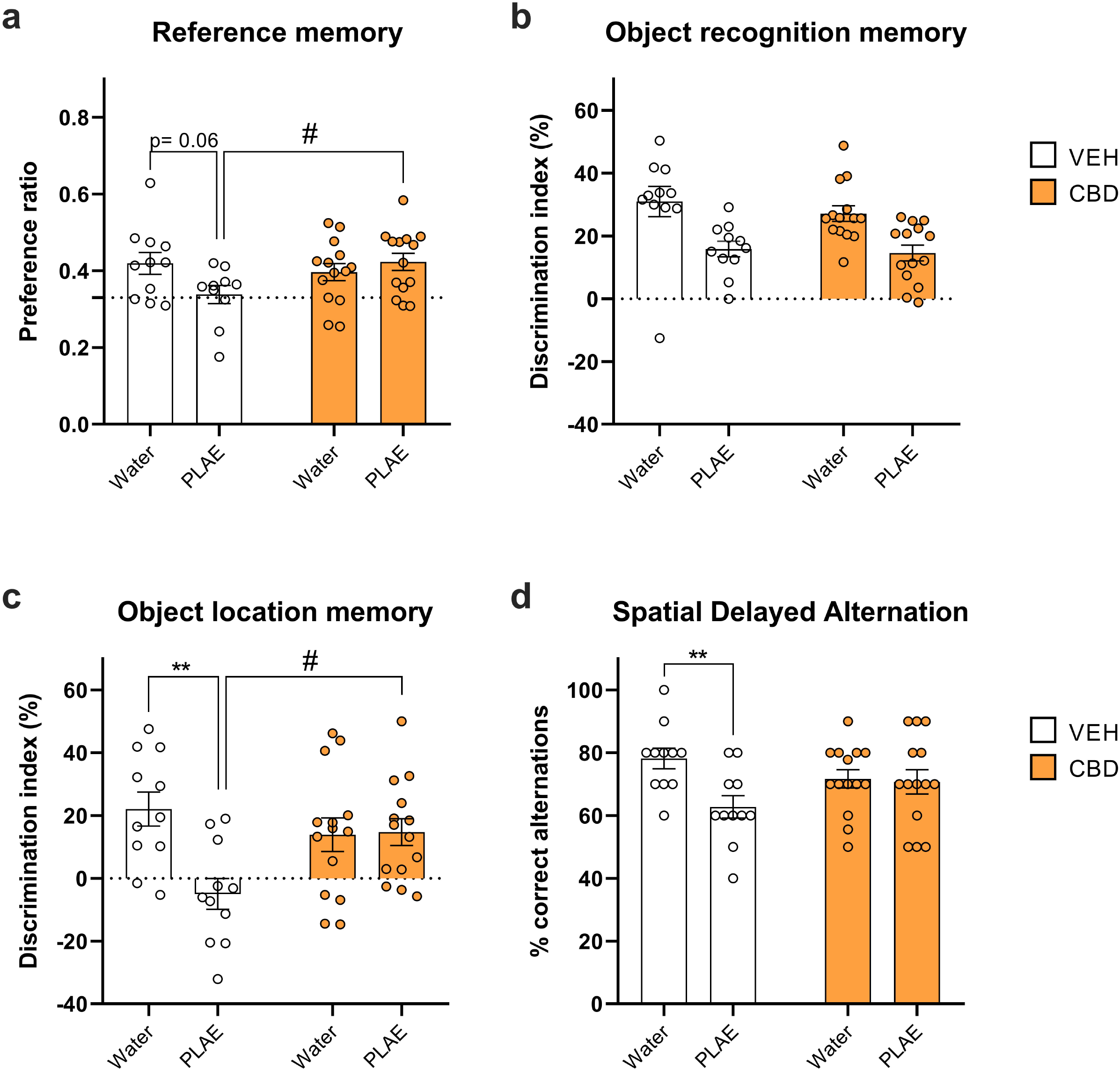
Protective effects of CBD on cognitive deficits induced by early alcohol exposure. (a) Preference ratio in Y-maze as reference memory evaluation (n=10-14 per group). The dashed grid line indicates the random preference that would be found if any animal spent an equal time in each of the three arms. (b) Percentage of discrimination index in the NOR task to assess object recognition memory (n=11-14 per group). (c) Percentage of discrimination index in the NOL task to test object location memory (n=11-14 per group). (d) Percentage of correct alternation in Y-maze as spatial working memory assessment (n=11-14 per group). Data are presented as mean ± SEM. Bonferroni’s post-hoc test **p<0.01 PLAE-VEH vs Water-VEH; #p<0.05 PLAE-CBD vs PLAE-VEH. CBD, cannabidiol; PLAE, prenatal and lactation alcohol exposure; VEH, vehicle

#### 3.2.2. CBD does not affect the recognition memory

Two-way ANOVA analysis of the percentage of discrimination index between novel and familiar object exhibits *group* effect (F_(1,46)_ = 19.98, p<0.001), without *treatment* or *interaction* effect (Figure 2b). Clearly, the PLAE mice showed a lower discrimination index, indicating an impaired recognition memory, regardless of the treatment.

#### 3.2.3. CBD ameliorates the PLAE-induced dysfunction in object location memory

The two-way ANOVA reveals a *group* (F_(1,46)_ = 6.738, p<0.05) and *interaction* effect between *group x treatment* factors (F_(1,46)_ = 7.598, p<0.01). Bonferroni’s multiple comparisons unveil that PLAE-VEH group exhibited significantly lower discrimination index compared to Water-VEH mice (p<0.01). In addition, PLAE-CBD animals performed more efficiently the task compared to PLAE-VEH (p<0.05), showing greater discrimination index (Figure 2c). Again, revealing that CBD treatment impedes the deleterious PLAE-induced effect on object location memory.

#### 3.2.4. CBD faintly hinders spatial working memory deficits induced by PLAE

Two-way ANOVA analysis displays a *group* (F_(1,46)_ = 5.605, p<0.05) and *interaction* (F_(1,46)_ = 4.379, p<0.05) effects on the percentage of correct alternations in the spatial delayed alternation task (Figure 2d). Subsequent Bonferroni’s *post-hoc* analyses unveil that PLAE-VEH mice carried out lower correct alternations than Water-VEH group (p<0.01). Additionally, the PLAE-CBD group was not significantly different as compared to PLAE-VEH, but neither to Water-VEH. Thus, this result supports that CBD is able to partially prevent spatial working memory deficits caused by PLAE.

### 3.3. CBD exerts slight effects on long-term neuroinflammatory response induced by PLAE

Two-way ANOVA analysis of PFC reveals a *group* effect for TNFα (F_(1,4)_ = 5.314, p<0.05), IL-6 (F_(1,39)_ = 7.987, p<0.01), COX-2 (F_(1,39)_ = 4.665, p<0.05) and caspase-3 (F_(1,40)_ = 11.29, p<0.01), having increased levels of these molecules in PLAE group regardless of treatment (Figure 3a-f). A tendency towards an *interaction* effect between *group x treatment* factors was found for caspase-3 (F_(1,40)_ = 3.53, p=0.06). Furthermore, we analysed whether the altered inflammatory markers might predict the caspase-3 elevation, as an indication of detrimental neuroinflammation. Pearson’s correlation showed significant positive correlations between levels of caspase-3 and TNFα (r=0.671, p<0.001) and between caspase-3 and COX-2 (r=0.8293, p<0.001). Thus, TNFα and COX-2 in PFC might be useful to predict apoptotic cellular death. However, the Pearson’s correlation was not significant for correlation between levels of caspase-3 and IL-6 (r=0.2480, p=0.1046) (Figure 3g).

**Fig. 3.**
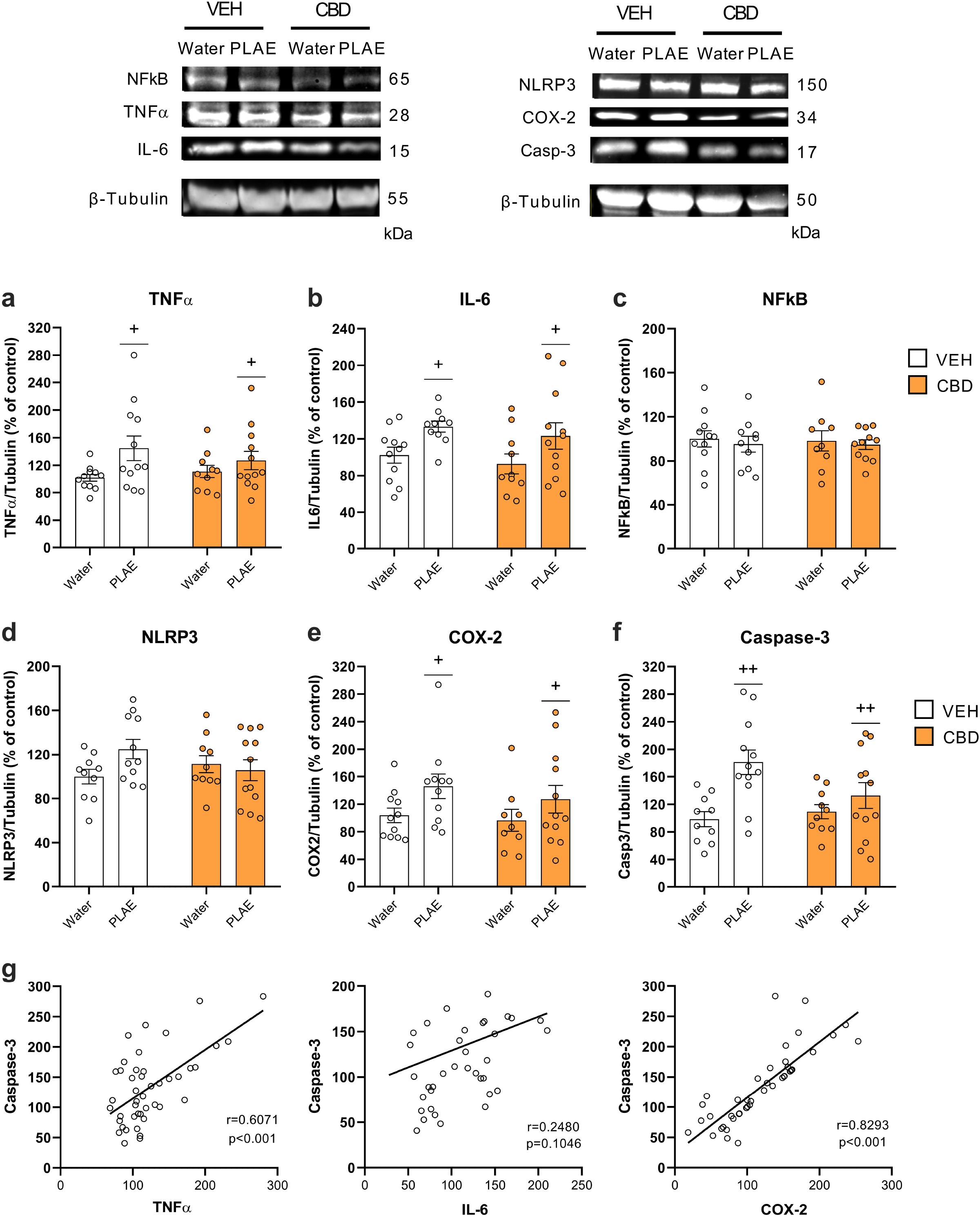
CBD effect on long-term pro-inflammatory response within prefrontal cortex induced by developmental alcohol exposure. (a-f) Western blot analyses of neuroinflammatory markers (TNFα, IL-6, NFkB, NLRP3, COX-2) and apoptotic marker (caspase-3) in PFC (n=9-12 per group). (g) Pearson’s correlation between levels of TNFα, IL-6 and COX-2 with Caspase-3. Data are presented as mean ± SEM. *Group* effect of two-way ANOVA test is represented by +p<0.05 and ++p<0.01 PLAE vs Water. PLAE-CBD vs PLAE-VEH. CBD, cannabidiol; Casp-3, caspase-3; COX-2, cyclooxygenase-2; IL-6, interleukin-6; NLRP3, NOD-like receptor protein 3; PLAE, prenatal and lactation alcohol exposure; TNFα, tumor necrosis factor alpha; VEH, vehicle

Two-way ANOVA of HPC unveils a *group* effect for TNFα (F_(1,41)_ = 18.35, p<0.001), NFkB (F_(1,39)_ = 6.375, p<0.05) and COX-2 (F_(1,40)_ = 4.857, p<0.05), showing that PLAE group had higher levels as compared to Water group. *Interaction* effects were found for TNFα (F_(1,41)_ = 4.302, p<0.05) and IL-6 (F_(1,36)_ = 4.343, p<0.05). Bonferroni’s multiple comparisons demonstrate that PLAE-VEH group had greater levels of TNFα (p<0.001) and IL-6 (p<0.05) than Water-VEH group. In addition, PLAE-CBD group showed lower levels of TNFα than PLAE-VEH group (Figure 4a-f), indicating somehow a beneficial effect of CBD treatment. In case of HPC we did not analyse correlations because the caspase-3 protein was not modified by PLAE.

**Fig. 4.**
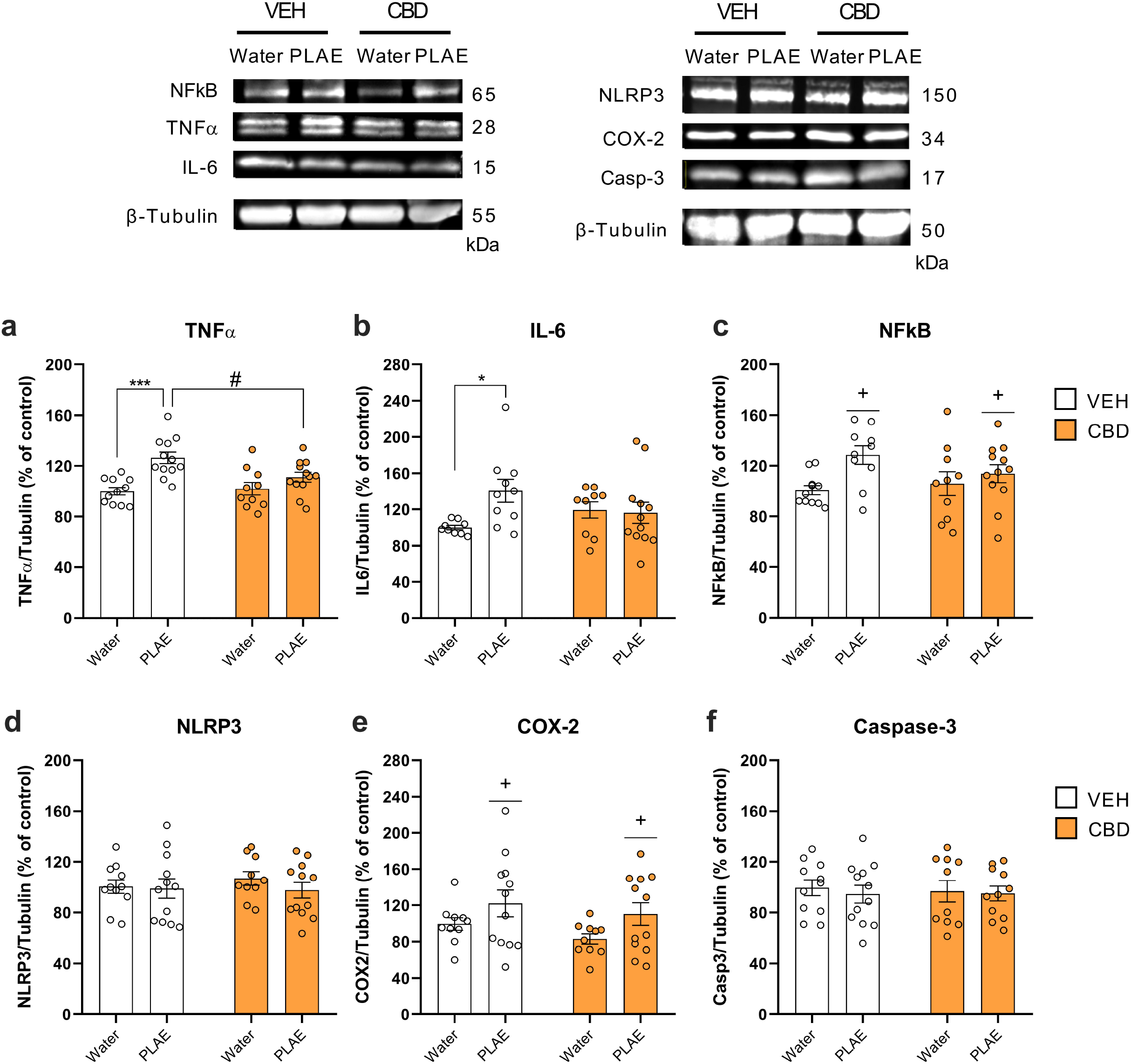
CBD effect on long-term pro-inflammatory response within hippocampus induced by developmental alcohol exposure. (a-f) Western blot analyses of neuroinflammatory and apoptotic markers in HPC (n=9-12 per group). Data are presented as mean ± SEM. *Group* effect of two-way ANOVA test is represented by +p<0.05 and ++p<0.01 PLAE vs Water. Bonferroni’s post-hoc test *p<0.05 and ***p<0.001 PLAE-VEH vs Water-VEH; #p<0.05 PLAE-CBD vs PLAE-VEH. CBD, cannabidiol; Casp-3, caspase-3; COX-2, cyclooxygenase-2; IL-6, interleukin-6; NLRP3, NOD-like receptor protein 3; PLAE, prenatal and lactation alcohol exposure; TNFα, tumor necrosis factor alpha; VEH, vehicle

## 4. DISCUSSION

In the present study, we demonstrate that CBD administered sub-chronically throughout a peri-adolescence period can counteract cognitive impairments induced by PLAE in male and female mice. Furthermore, our molecular findings confirm that PLAE long-term neuroinflammatory state in the PFC and HPC can be attenuated by CBD treatment.

The CNS is especially vulnerable to the alcohol effects, primarily throughout developmental periods, like pregnancy and lactation [37]. FASD’s most severe manifestations involve behaviour and learning difficulties [3]. In fact, clinical reports have documented marked deficits in high-order functions, such as working memory [38], spatial memory [39,40], and delayed object memory [41]. Our FASD-like mouse model partly resembles the neurobehavioural outcomes reported in those clinical studies, since PLAE animals exhibit dysfunctions on working memory, spatial memory, as well as object recognition and object location memory.

Although the behavioural consequences of alcohol exposure during gestation and lactation periods are well characterized, the underlying molecular mechanisms are still poorly understood. In this sense, preclinical models provide the opportunity to gain knowledge on those potential mechanisms responsible for FASD. Among the putative mechanisms, it is well described the glia activation following moderate-to-high levels of alcohol exposure in rodent models of FASD [6,9]. This activation is accompanied by increased levels of pro-inflammatory cytokines in brain areas, including the cerebral cortex and the HPC [6,13,42], which could remain elevated until the adulthood [10,43]. In line with these findings, we show a long-term neuroinflammatory response within PFC and HPC induced by gestational and lactation alcohol exposure. PLAE mice exhibit higher levels of TNFα, IL-6 and COX-2 in both PFC and HPC.

Neuroinflammatory processes could be either neuroprotective or cytotoxic, promoting tissue homeostasis or enhancing tissue damage [44]. TLR4 is considered a major contributor to innate immune response and a key mediator to activate neuroinflammation following alcohol exposure [12,45]. Moreover, evidence suggests that alcohol-induced apoptotic signalling is mediated by TLR4 [12,46]. In addition, prior studies stated cell degeneration in cerebellar vermis while no signal of neuronal loss was found in HPC, despite reporting an increased pro-inflammatory cytokines profile within both regions due to neonatal alcohol exposure [47]. Similarly, we have displayed that PLAE induces long-term brain region-dependent effects on apoptotic signalling, given that caspase-3 levels were higher in PFC, while no changes were found in HPC, as previously reported [10]. Moreover, increased levels of both TNFα and COX-2 positively correlate to caspase-3 elevation in PFC, supporting the involvement of pro-inflammatory response in triggering the apoptotic signalling in this brain area. In contrast, PLAE increases NF-kB in HPC without modifications in PFC. Emerging studies support a pivotal role for NF-kB in the pathogenesis of many neurological disorders, either enhancing or mitigating disease [48]. NF-kB mediates neuroinflammatory response and neurodegeneration following excessive and acute alcohol exposure, albeit adaptive changes in NF-kB signalling were suggested to protect brain cells against potent neurotoxicity after repeated cycles of alcohol consumption and withdrawal [49]. In our model, mice were exposed to alcohol repeatedly throughout six weeks, but the brain extractions were carried out after a period of withdrawal. Therefore, the increased levels of NF-kB might enhance anti-apoptotic pathways in HPC and consequently no caspase-3 deregulations are reported in this area. On the other hand, the pro-apoptotic caspase-3 elevation in PFC could be due to this region dependent lack of NF-kB activation.

However, other concomitant mechanisms occurring within the HPC, like impaired neurogenesis [27] cannot be discarded and should be further evaluated.

The role of neuroinflammation as one of the underlying causes of the behavioural deficits induced by early alcohol exposure is further supported by genetic and pharmacological research. Mice lacking TLR4 do not show behavioural impairments or changes in pro-inflammatory markers following prenatal alcohol exposure indicating that the FASD phenotype is at least partially TLR4 dependent [12]. In the same way, anti-inflammatory drugs like ibuprofen [50] and resveratrol [13] have been proven useful to lessen the FASD phenotype.

Cannabinoids represent an alternative anti-inflammatory strategy to treat several diseases [14,51]. Indeed, they might exert positive or negative effects on squeals caused by excessive alcohol use, depending on the cannabinoid compound [15]. Although the detrimental consequences of CB1R agonists on developing brain have been well established [52,53], CBD has recently emerged as promising candidate for several pathological conditions, including alcohol-related harms [54]. Given that preliminary clinical data has already shed light on the positive CBD effects on FASD [55,56], preclinical evidence is firmly necessary. Here, we demonstrate that CBD treatment during peri-adolescence period can attenuate cognitive deficits induced by PLAE, without modifications in control group. Specifically, reference and object location memories were improved by CBD. An interaction between factors was also found for delayed spatial working memory, showing that CBD might slightly attenuate PLAE-induced working memory defect. On the other hand, CBD does not rescue recognition memory deficits. Therefore, the main positive effect of CBD’s is on spatial memory. Similarly, other studies conducted to evaluate neurological disorders proposed that CBD shows therapeutic benefit for spatial and working memory deficits induced by hepatic encephalopathy [57], hypoxic brain injury [58], as well as in preclinical models of infection-induced inflammatory disorders like polyinosinic-polycytidilic acid infection [59] or cerebral malaria [20]. Notwithstanding, other reports propose a dose-dependent CBD effect on recognition memory improvement. For instance, higher doses of CBD counteracted recognition memory impairment induced by a pharmacological model of schizophrenia, whereas lower doses did not [60]. Overall, CBD administration appears to improve cognitive deficits in several domains, which may vary depending on the pathological condition and/or the dose of CBD.

Attenuation of pro-inflammatory signalling following CBD administration was associated with cognitive improvements in several preclinical models [58,61]. In the present study, we observe a clear long-term reduction of hippocampal TNFα in PLAE mice treated with CBD. Down-regulated expression of hippocampal TNFα and/or its receptor 1 has been associated with improved spatial and working memory after CBD treatment [57,62]. Conversely, another study has associated CBD-induced improvement of associative learning with a down-regulation of TNFα expression in frontal cortex, but not in HPC [63]. Moreover, we exhibit a slight CBD effect to counteract IL-6 within the HPC. On the opposite, our findings show no long-term CBD anti-inflammatory effects on PFC. Hence, CBD might be exerting brain region-dependent long-lasting anti-inflammatory mechanisms. Previously, others have displayed that CBD-induced anti-inflammatory response might be region dependent [63]. Besides, CBD showed effective neuroprotection in the HPC at 30mg/kg dose, while no significant results were demonstrated at 3 mg/kg in a mouse model of brain ischemia [58]. In this sense, future research focusing on elucidating the brain-region specific long-lasting inflammatory modulation induced by CBD would be crucial to ameliorate FASD phenotype.

Overall, CBD appears to be a promising therapeutic drug since it could hamper cognitive impairments caused by several pathological conditions, potentially through neuroimmune modulation. In regards to FASD, this is the first preclinical study exhibiting the beneficial effects of CBD on PLAE-induced cognitive deficits, along with long-lasting reduction of pro-inflammatory cytokines, specifically within HPC. Thus, this study provides new insights into the role of non-psychotropic cannabinoids as pharmacological strategy to counteract alterations in cognitive performance of FASD. However, further studies evaluating the dosage-dependent positive or adverse effects of CBD would be necessary to finally elucidate the most effective period and conditions to administer CBD.

## AUTHORS’ CONTRIBUTIONS

Alba Garcia-Baos and Olga Valverde were responsible for the study concept and design. Alba Garcia-Baos and Xavi Puig-Reyne carried out the experimental studies. Alba Garcia-Baos, Olga Valverde and Óscar García-Algar participated in the interpretation of findings. Alba Garcia-Baos and Olga Valverde drafted the manuscript. All authors critically reviewed the content and approved the final version for publication.

## ACKNOWLEDGEMENTS

This study was supported by the Ministerio de Economia y Competitividad (grant number PID2019-104077RB-100), Ministerio de Sanidad (Retic-ISCIII, RD16/017/010 and Plan Nacional sobre Drogas 2018/007). Alba Garcia-Baos received a FI-AGAUR grant from the Generalitat de Catalunya (2019FI_B0081). The authors wish to thank Laia Alegre Zurano for manuscript proofreading. We also thank Phytoplant Research S.L. for CBD supply.

## ABBREVIATIONS

CBD: cannabidiol
CNS: central nervous system
COX: cyclooxygenase
CB1R: cannabinoid receptor type 1
CB2R: cannabinoid receptor type 2
DID: drinking in the dark
FAAH: fatty acid amide hydrolase enzyme
FASD: Foetal alcohol spectrum disorder
HPC: hippocampus
IL: interleukin
NF-kB: nuclear factor Kappa-B
NLRP: NOD-like receptor protein
NOL: novel object location
NOR: novel object recognition
PD: post-natal day
PFC: prefrontal cortex
PLAE: prenatal and lactation alcohol exposure
PPAR: peroxisome proliferator-activated receptor
TNFα: tumor necrosis factor alpha
TLR4: toll-like receptor 4
VEH: vehicle.

